# Comparative genome analysis of *Corynebacterium ulcerans* Japanese isolates revealed domestically and globally diverse geographical distribution of the organism of different types

**DOI:** 10.64898/2026.04.05.716610

**Authors:** Miyuki Kimura, Tsuyoshi Sekizuka, Makoto Kuroda, Mitsutoshi Senoh, Motohide Takahashi, Akihiko Yamamoto, Masaaki Iwaki

## Abstract

*Corynebacterium ulcerans* is a zoonotic pathogen causing infections in humans that are clinically indistinguishable from those of *Corynebacterium diphtheriae*. In many developed countries, companion animals such as dogs and cats, are the predominant sources of infection by *C. ulcerans,* as is the case in Japan. To date, we have collected 106 Japanese clinical and domestic animal isolates of *C. ulcerans* and a recently branched species, *C. ramonii*, most of which were isolated from humans and cats. In the present study, we performed a comparative analysis of whole-genome sequences from Japanese isolates and 597 published isolates from Japan and outside Japan. The Japanese isolates were divided into two distinct lineages: *C. ulcerans* and *C. ramonii*. The MLST types of *C. ulcerans* Japanese isolates exhibited a unique distribution pattern, with one major type (ST337) accounting for 69.0%, which is very rare in Europe. In addition to Japan, distinct MLST pattern compositions were observed across geographically distinct regions. The sequence types were associated with lysogenized prophage types that encode the diphtheria toxin (*tox*) gene and were partially associated with toxin production under certain conditions. Through SNV analysis, transmission from animals to humans has been suggested in some clinical cases.

**Significance Statement:** By analyzing more than 700 genomes, we demonstrate striking geographic differences in multilocus sequence types and toxigenic prophage compositions between Japanese and European isolates. Notably, we identify the predominance of sequence type ST337 in Japan, a lineage that is rare in Europe, and show that prophage types encoding the diphtheria toxin are closely associated with specific sequence types and toxin production phenotypes. Our single-nucleotide variant analyses further provide genomic evidence supporting zoonotic transmission between companion animals and humans in several clinical cases.

## Introduction

*Corynebacterium ulcerans* is a zoonotic pathogen that causes diphtheria-like infections in humans, because of which, its clinical manifestations remain indistinguishable from those of *Corynebacterium diphtheriae*. Thus, the World Health Organization categorizes *C. ulcerans* infection into "diphtheria" [1, 2]. In recent years, the number of *C. ulcerans* infection cases has increased in the developed countries, and in Japan, the number of these cases exceeds that of *C. diphtheriae* infection [1–3]. In Japan, no diphtheria cases caused by *C. diphtheriae* have been observed since the last case in 1999, whereas 34 human *C. ulcerans* cases have been reported from 2001 to 2019, with two fatal cases [3, 4].

Historically, *C. ulcerans* has been known to cause accidental infection due to drinking unpasteurized milk from cows with mastitis caused by *C. ulcerans* [5, 6]. However, most human *C. ulcerans* infections have recently been associated with companion animals such as cats and dogs [1, 2, 7, 8], which is also the case in Japan [3]. In most of the 34 Japanese human cases, including two fatal cases, the presence of accompanying companion animals, mostly cats, has been demonstrated [3, 4]. In addition, *C. ulcerans* has been isolated from dogs, including custodial and hunting dogs [9–11]

The host range of *C. ulcerans* extends to livestock and various wildlife [12], including heifers [13], wild boars and roe deer [14, 15], hedgehogs [16, 17], squirrels [18], shrew moles and owls [19], killer whales and lions in zoos [20], and macaque monkeys as experimental animals [21]. This wide host range of *C. ulcerans* contrasts with the narrow range of *C. diphtheriae,* found exclusively in humans, with very limited exceptions in horses [12, 22, 23].

The most important virulence factor *in C. ulcerans* is diphtheria toxin (also called diphtheria-like toxin). The toxin structurally resembles (∼95% homology at amino acid and nucleotide levels) to that produced by *C. diphtheriae* [8, 24]. In the case of *C. diphtheriae*, the 58 kDa toxin is one of the most intensively studied bacterial toxins [8]. The toxin binds to the target cell surface receptor HB-EGF, gets internalized into the lysosome, released in the cytosol by lysosome acidification, and is ultimately modified by ADP-ribosylation of one of the ribosome components, EF-2, targeting its diphthamide residue, resulting in inactivation of ribosomes and killing the cell [8] [25, 26]. The expression of the toxin (*tox*) gene is repressed by the dtxR product [27] and deregulated in the absence of iron in the environment [28]. Similar to *C. diphtheriae* [29, 30], the *tox* in *C. ulcerans* is usually located in the prophage genome and is lysogenized in the host bacterial genome. It is generally agreed that toxigenicity is mediated by the shared *tox*-bearing bacteriophages between *C. ulcerans* and *C. diphtheriae* by shared *tox*-bearing bacteriophages [31–34]. In 2012, Sekizuka et al. [35] demonstrated using whole-genome sequencing of the *C. ulcerans* 0102 strain that the *tox*-bearing prophage in the genome of this strain was distinct from known *tox*-bearing *C. diphtheriae* prophages, with no significant homology in any structural genes except *tox*. This finding is supported by previous and subsequent studies using genome sequencing of European isolates [36–38]. In particular, a recent analysis by Crestani et al. of 582 isolates, most of which originated in France and Belgium, showed the presence of multiple types of prophages across several typical lineages [39].

Ribotyping has been the molecular typing method of *C. ulcerans* as the gold standard for years, adopting the method for *C. diphtheriae* [40, 41] and *C. ulcerans* [42]. Subsequently, discriminative multilocus sequence typing (MLST) [43] has become dominant and widely used, gradually being substituted for core-genome MLST (cgMLST) [16, 39]. To date, conventional MLST data have been compiled in databases and are publicly accessible (https://bigsdb.pasteur.fr/diphtheria/). To date, we have sequenced 106 Japanese *C. ulcerans* isolates from humans and non-human hosts, mainly companion animals. In the present study, using genome analysis of these isolates and previously published isolates, a unique distribution pattern in conventional MLST and prophage types was demonstrated in relation to toxin production. Case analyses of single-nucleotide variants (SNV) in human cases accompanied by companion animals are also presented.

## Results

### 1. Lineages and geographical distribution

The genomes of 106 Japanese *C. ulcerans* isolates were sequenced and analyzed alongside publicly available genome sequences of Japanese and European isolates. The analyzed sample consisted of 122 Japanese isolates: humans (38), cats (70), dogs (8), macaque monkey (1), killer whale (1), cat dwellings (3), and unknown origin (1). In 2025, Crestani et al. [39] published a large-scale analysis of 587 genome sequences, most of which originated in France and Belgium. Our analysis included a previously published dataset of sequences [39]. Figure 1 shows the high-resolution core-genome phylogeny of the available genome sequences (703 in total), including those of some *C. ramonii* isolates. The two major clades shown in Figure 1 represent *C. ulcerans* and *C. ramonii*. As shown in the Figure, a large portion of Japanese *C. ulcerans* isolates were clustered (lower part of circle 3) compared with European isolates, which were distributed across a wide range of clades. Host-animal- or isolation-year-specific clades were not identified in the phylogenetic analysis. As expected, MLST and the presence of *tox* were closely related to clades. Comparison of the MLST proportions of Japanese isolates with those of French and Belgian isolates revealed a global geographical difference in MLST proportions (Figure 2). With the information obtained by conventional MLST analysis of the Japanese *C. ulcerans* genome sequences obtained in the present study and those available from public databases in October 2025 (100 in total), ST 337 was shown to be the predominant sequence type, accounting for 69.0 % (69 isolates out of 100. Figure 3, see also left panel and Figure S1, shown in light blue color in column 4 in the lower clade), followed by ST 325 (12 isolates, 12.0%). In addition to *C. ulcerans*, 25 ST344 strains representing *C. ramonii* [63] were found among the isolates (not shown in Figure 3). French isolates were more diverse than Japanese isolates and comprised more STs. The Japanese majority ST 337 comprised only 2.2% (10 isolates); the major STs in France were ST 325 (170 isolates, 38.2%), ST 339 (54 isolates, 12.1%), and ST 331 (70 isolates, 15.7%). Among the Belgian isolates, the majority were ST 331 and ST 332 ( 15 isolates, 36.6% each). In Germany, ST331 was the most prominent, followed by STs 325, 326, and 332. In the United Kingdom (data from the MLST site of the Institut Pasteur (https://bigsdb.pasteur.fr/diphtheria/)), ST 325 was not observed, and ST 543, which is rare in France, was the most common.

**Figure 1.**
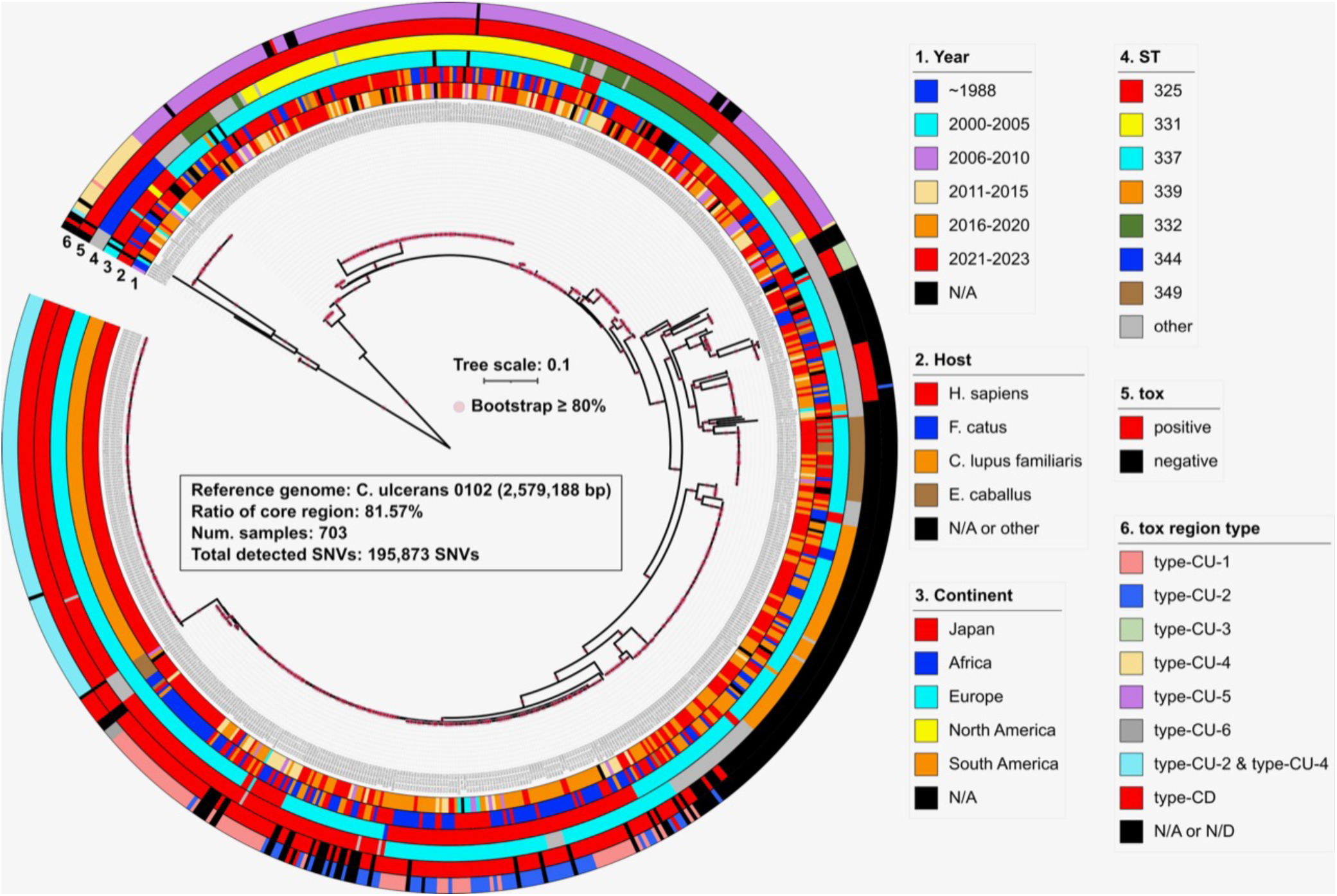
Core-genome phylogeny and metadata of genomes of 703 C. ulcerans and C. ramonii isolates. Seven hundred three genome sequences available thus far, including those obtained in the present study, were analyzed as described in the Materials and Methods section, and expressed in a phylogenetic tree. Metadata associated with each isolate, as well as that obtained from the database, is also shown in the boxes. The circles in the Figure represent 1, year of isolation; 2, host animal; 3, place of isolation (continent or country); 4, sequence type (MLST); 5, presence/absence of *tox* gene; and 6, *tox* region type.

**Figure 2.**
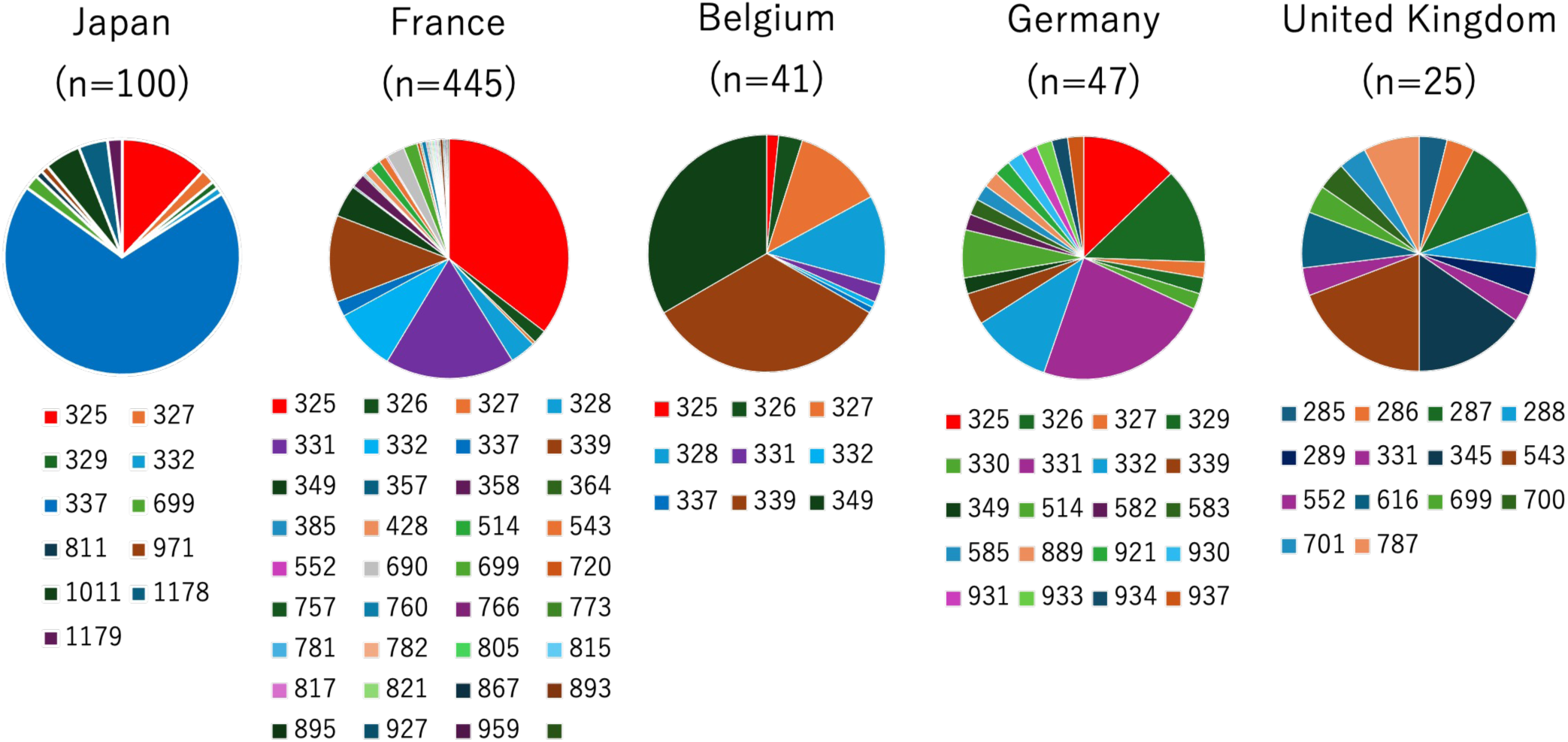
C. ulcerans MLST types in Japan and European countries. MLST types of *C. ulcerans* isolates and those extracted from genomic sequences available in November 2025, except for those of *C. ramonii* (e.g., ST344), are illustrated. The proportion of STs in each country is indicated by circles. The STs are indicated by the colors shown below each circle.

**Figure 3.**
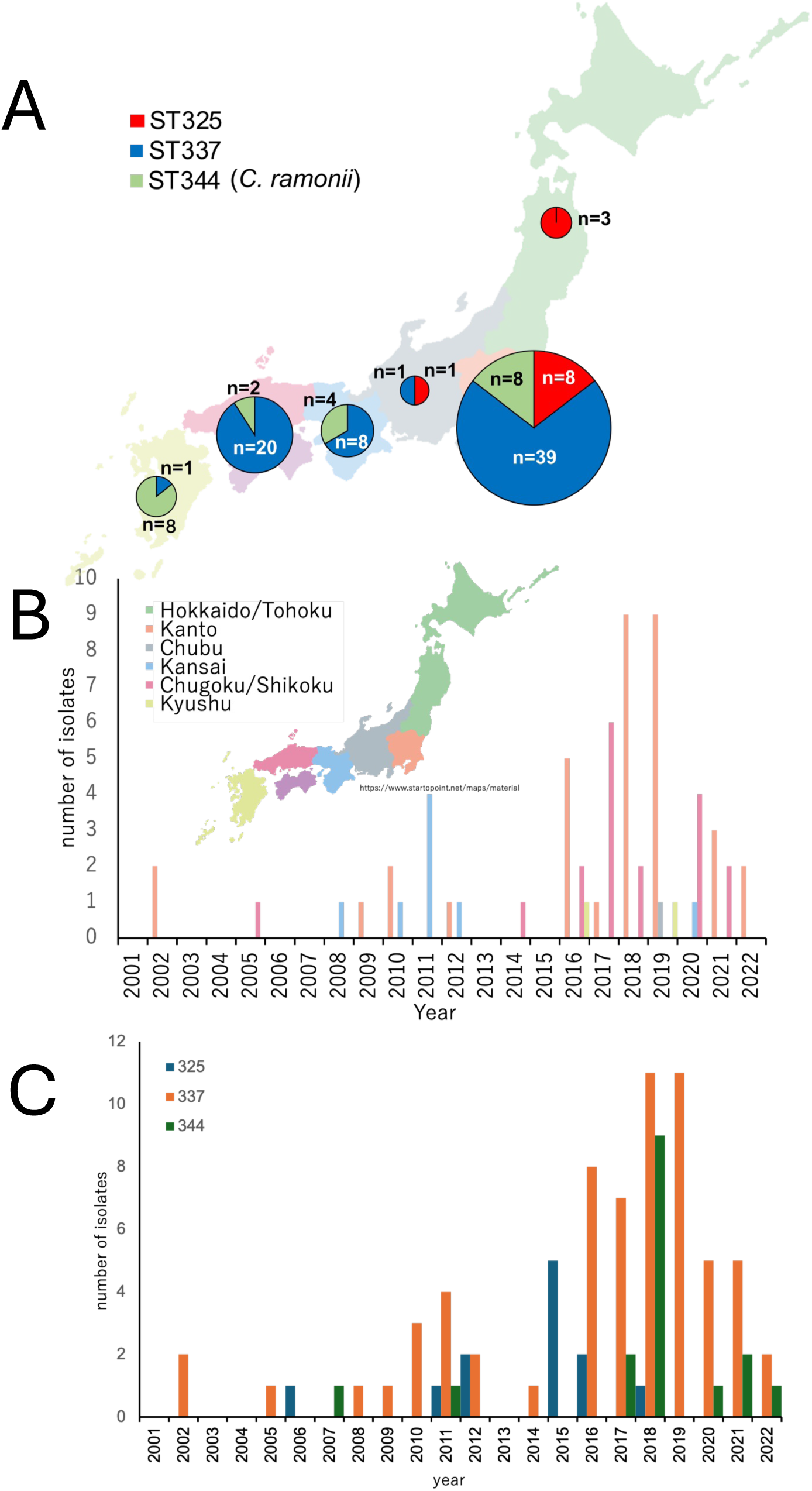
C. ulcerans and C. ramonii in Japan. **A.** Domestic distribution of major STs of *C. ulcerans* and *C. ramoni*i in Japan. Distribution of isolates belonging to two major MLST types (ST337 and ST325), as well as that of *C. ramonii* (ST344), is shown. The land of Japan is divided into six isolated areas (from north to south: Tohoku/Hokkaido, Kanto, Chubu, Kansai, Chugoku/Shikoku, and Kyushu), and the proportion of isolates in each area is shown in circles. **B.** Geographic transition of *C. ulcerans* ST337 isolates in Japan from a chronological perspective.The number of ST337 isolates is reported per year across six regions of Japan. The regional scheme is the same as in Figure 3: Tohoku/Hokkaido, Kanto, Chubu, Kansai, Chugoku/Shikoku, and Kyushu, from north to south. **C.** Chronological representation of the isolates in Japan.The numbers of the two major STs of *C. ulcerans* (ST337 and ST325), as well as that of *C. ramonii* (ST344), are displayed by year. Bars represent each ST.

The distributions of major STs in Japan are shown in Figure 3A. ST337 (blue) isolates were distributed from the center to the west. In contrast, isolation of ST 325 (red) was limited to the middle to northern regions of the country. In the northern region, all isolates were ST325. In addition, the distribution of ST344 (green: C. ramonii) shifted further west, comprising more than 80% of isolates from the westmost region. All three STs were detected in the central region (including the capital city of Tokyo), approximately reflecting the proportion of each ST among the total isolates in the country. A chronological view of the three major STs is presented in Figure 3B. Isolation began in 2001 (ST337) and continued at a low level until 2010, when the first peak began to increase and was completed in 2012. After the blank period, the second peak starts in 2014 and diminishes in 2022. All STs exhibited similar behavior. For geographic distribution per animal of origin and prophage type, see Supplementary Figures S2 and S3, respectively.

Further analysis was performed on the dominant ST337 strain (Figure 3C). The first detection occurred in 2001 in the Kanto region of central Japan. The first peak around 2011 was predominantly in the Kansai region in the midwest of the country. After the disappearance of the peak, a geographical shift apparently occurred; that is, the largest peak later in 2021-2022 was not in the Kansai region but mainly in the Kanto region.

### 2. Prophages and attachment sites

Among the Japanese isolates, approximately 90% were toxigenic (harboring the *tox* gene and producing active toxin). In *C. ulcerans*, the *tox* gene is generally found on the chromosome as part of a prophage. In the present study, genome analysis revealed that five types of prophages exist among Japanese toxigenic *C. ulcerans* isolates (Figure 4), and in many cases, these types are associated with MLST types. The CU-1 and CU-2 prophage types were recognized as intact prophages containing genes and open reading frames essential for generating mature phage particles. PHASTEST analysis [64] revealed that the CU-1 and CU-2 prophage types are intact, harboring the genes required for mature phage particle assembly, and exhibiting high homology with PHAGE_Gordon_Nyceirae_NC_031004(10). Toxigenic ST337 isolates from Japan were exclusively lysogenized by these two types of prophages (Figure S1). The identical nucleotide sequences of the surrounding regions (*hisC*, *nagE*, *pdxT,* and other CDSs) showed that these two types of prophages integrated into the *C. ulcerans* chromosome in the same direction at attachment sites close to each other. Two other prophage types (CU-3 and CU-5) lacked most phage components, except for the *tox* gene. These types have the *tox* gene but retain only a small part of the bacteriophage components; thus, they could also be regarded as pathogenicity islands (PAIs), as reported by Meinel et al. [38] and Crestani et al. [39]. The common features of these incomplete and complete types (CU-1 and CU-2) are the surrounding regions. The high homology of the surrounding regions in all prophage types suggests that they share only a few common integration sites on chromosomes. The CU-4 type, which also represents an incomplete prophage, was exclusively associated with ST344 (*C. ramonii*).

**Figure 4.**
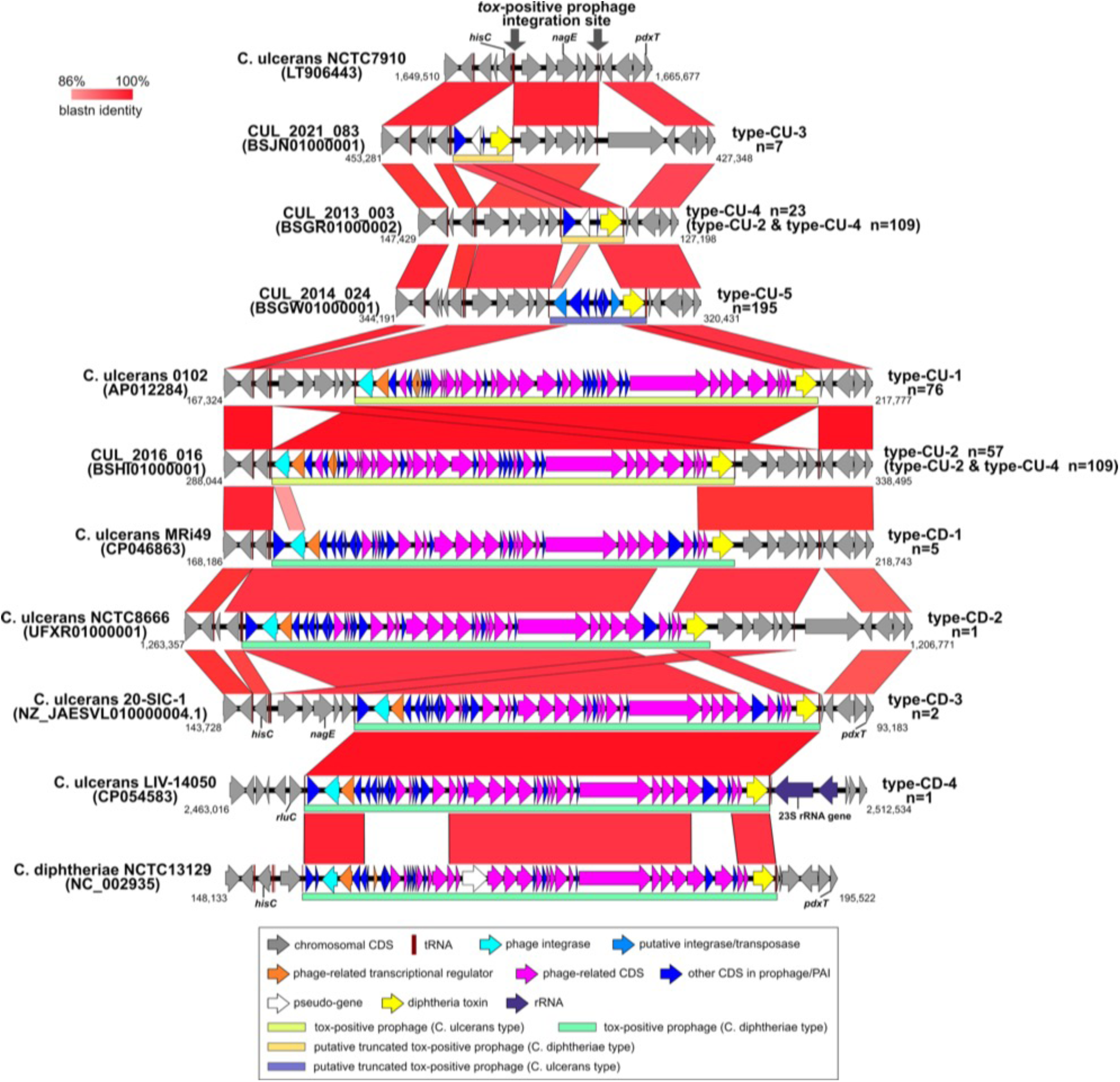
Schematic representation of prophage types. Prophage types determined by the analysis of genomic sequences of isolates and those obtained from databases are shown. Red squares/regions between the sequences indicate high homology. The other features are shown in the box below.

Some isolates from outside of Japan have different prophage types. They had a similar composition to the prophages lysogenized in *C. diphtheriae*. A typical example of *C. diphtheriae*-type prophage is the NCTC13129 strain shown in the lowermost part of Fig. 4. The nucleotide sequences of prophages of type CD-1-4 were highly homologous to those of NCTC13129. In contrast, no homology was observed between prophages of the type CD-1 (*C. diphtheiae* type) and CU-2 (*C. ulcerans* type) (Fig. 4, middle part), indicating that the CU-type prophages are different from CD-type prophages, although these prophages have a similar composition and order of components, as shown in Fig. 4. Interestingly, our extended analysis showed that two more types of prophages (Fig. 5, CU-6 and CD-5) were found in the isolates published by Crestani et al. [39]. The CU-6 type represents a “hybrid-type” of prophage of the CU- and CD- types. Another type of prophage, CD-5, was partially homologous to *C. diphtheriae* prophage (CD1036), but the remaining components did not show homology with any of the CU or CD types.

**Figure 5.**
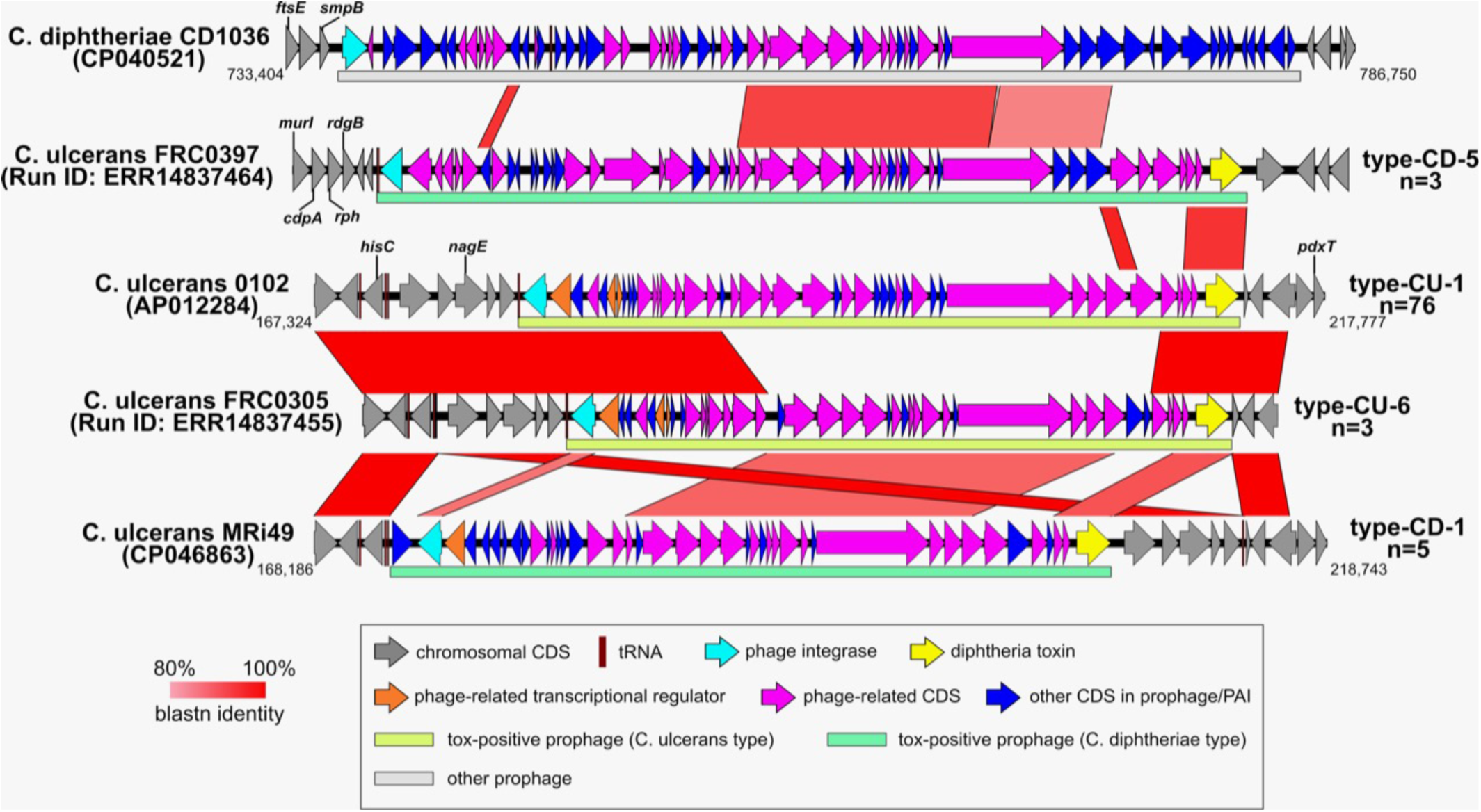
Schematic representation of atypical types of prophage regions. Atypical (hybrid) regions containing the *tox* gene are shown. Red squares/regions between the sequences indicate high homology. The other features are shown in the box below.

### 3. Case analyses

The listed *C. ulcerans* isolates comprised nine groups of clinical clusters isolated from human cases and accompanying cats (Fig. 6A). Cases 1–4 were clearly unrelated, with cases 3 and 4 possibly having some similarities. Cases 5–9 are closer to each other than the above cases. Geographically, Case 6 was from Okayama, which will be reported in a separate paper; Case 7 was from Tochigi; Case 8 was from Gunma; and Case 9 was from Chiba (Figure 6B). As shown in Figures 6A and 6 B, many were composed of ST337 organisms. The ST325 clusters were concentrated in the central (Kanto) region, unlike the ST337 clusters, which were spread across a wide range of geographical regions. Interestingly, a rare sequence type, ST699, formed a human-dog cluster in the mid-west (Kansai) region. An ST344 (*C. ramonii*) cluster was found in the extreme western (Kyushu) region, in accordance with the distribution shown in Fig.3 (see also Fig. S4).

**Figure 6.**
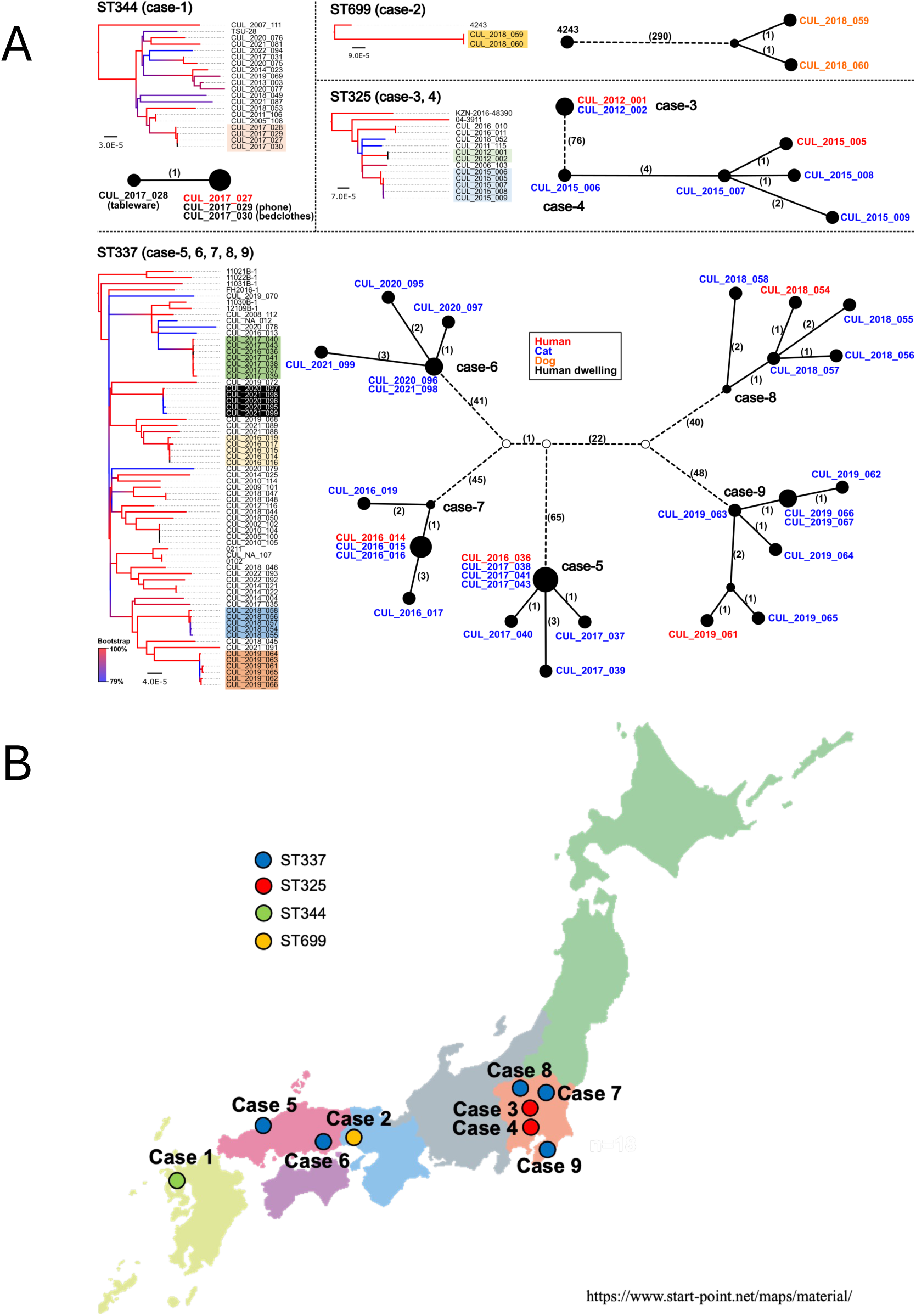
Case analyses. Nine human–cat and human-dog cluster cases were represented. **A**. SNV representations of the cases. The closed circles represent isolates with diameters corresponding to the same number of SNVs. Open circles represent putative nodes. The numbers on the lines represent the number of SNVs between the closed circles. **B**. Geographic representation of cases. Circles indicate the STs of the isolates in these cases.

### 4. Toxin production

Toxin production by the Japanese isolates was estimated by cultivating each isolate in Elek broth at a common optical density (∼0.5) and comparing the Vero cell cytotoxicity [44] of the culture filtrate. As shown in Figure 7, ST337 isolates demonstrated significantly lower toxin production (p<0.01) than STs 325, 332, and 344 isolates. Among the ST337 isolates, those that possess CU-2 type prophages showed significantly higher toxin production (p<0.01) than those that possess CU-1 type prophages. The ST344 isolates (*C. ramonii*) showed variable toxin production (Fig. 7). In particular, these two isolates exhibited very high levels of toxin production. The level of toxin production in these isolates was higher than that in the TSU-28 strain [65], which possesses two copies of the *tox* genes. Some outliers are observed in the ST325 dataset. However, no clear relationship between the nucleotide sequences of *tox*, the regulatory gene *dtxR,* and their respective promoter regions was observed when comparing the nucleotide sequences of individual isolates (Fig. 8A, color strips 1-3). Figure 8 B shows the nucleotide substitutions in the *tox* promoter regions in isolates with high-level ST344 producers (see below) and average-level producers. In high-level toxin-producing strains, two nucleotide substitutions were detected within an interrupted palindromic sequence of the *tox* promoter. In addition, pan-genome analysis revealed no accessory genes associated with high levels of toxin production (Fig S1).

**Figure 7.**
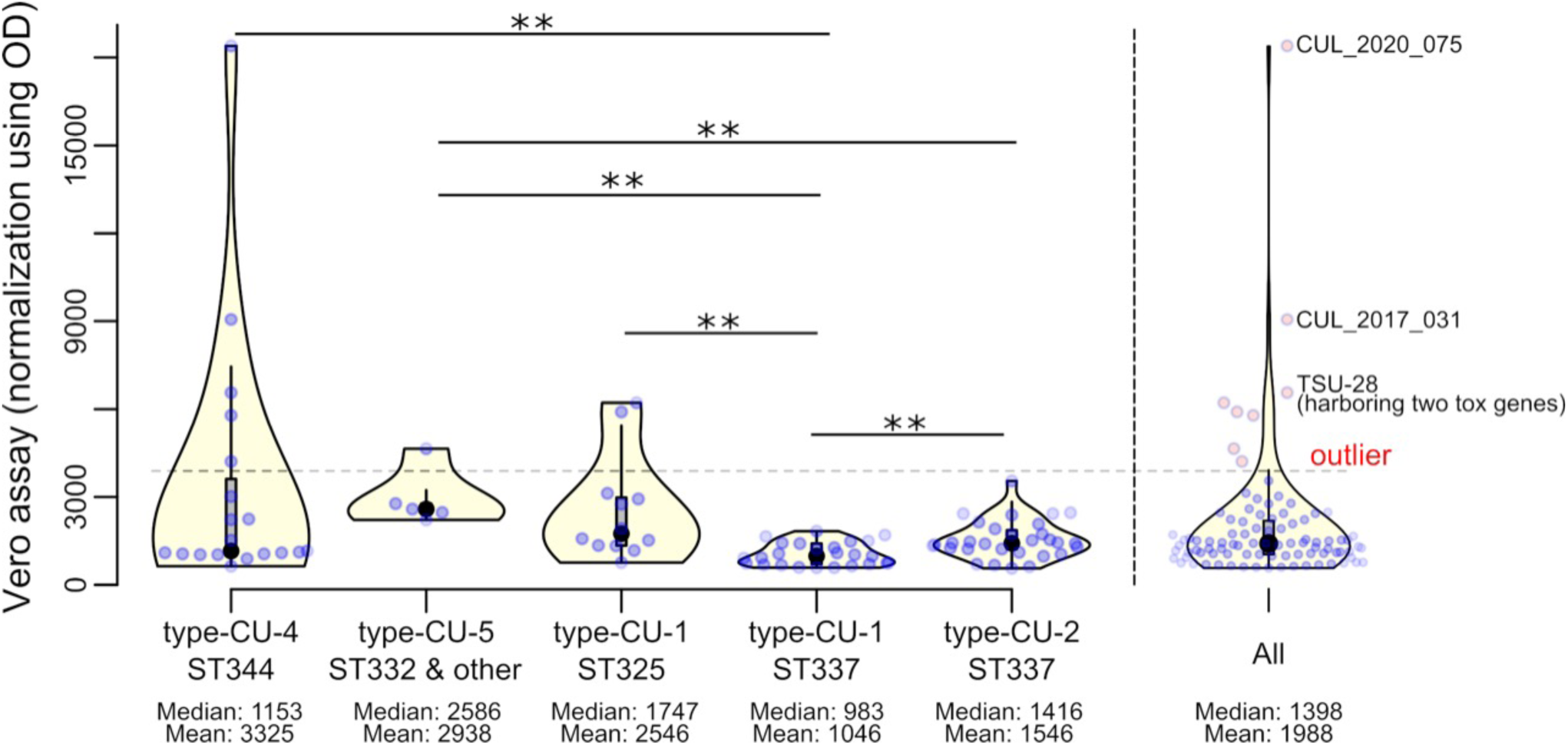
Toxin production by the isolates. Isolates were cultivated in Elek broth, and the toxin activity of the culture filtrate was measured by the Vero cell cytotoxicity assay, as described in the Materials and Methods. Levels of toxin production by MLST type (for ST337, further divided by prophage type) are shown in violin plots. Blue circles represent the toxin production level of each isolate, represented by the endpoint titer of the culture filtrate normalized to OD600=0.5, on the vertical axis. Asterisks (**) indicate significance at P<0.01.

**Figure 8.**
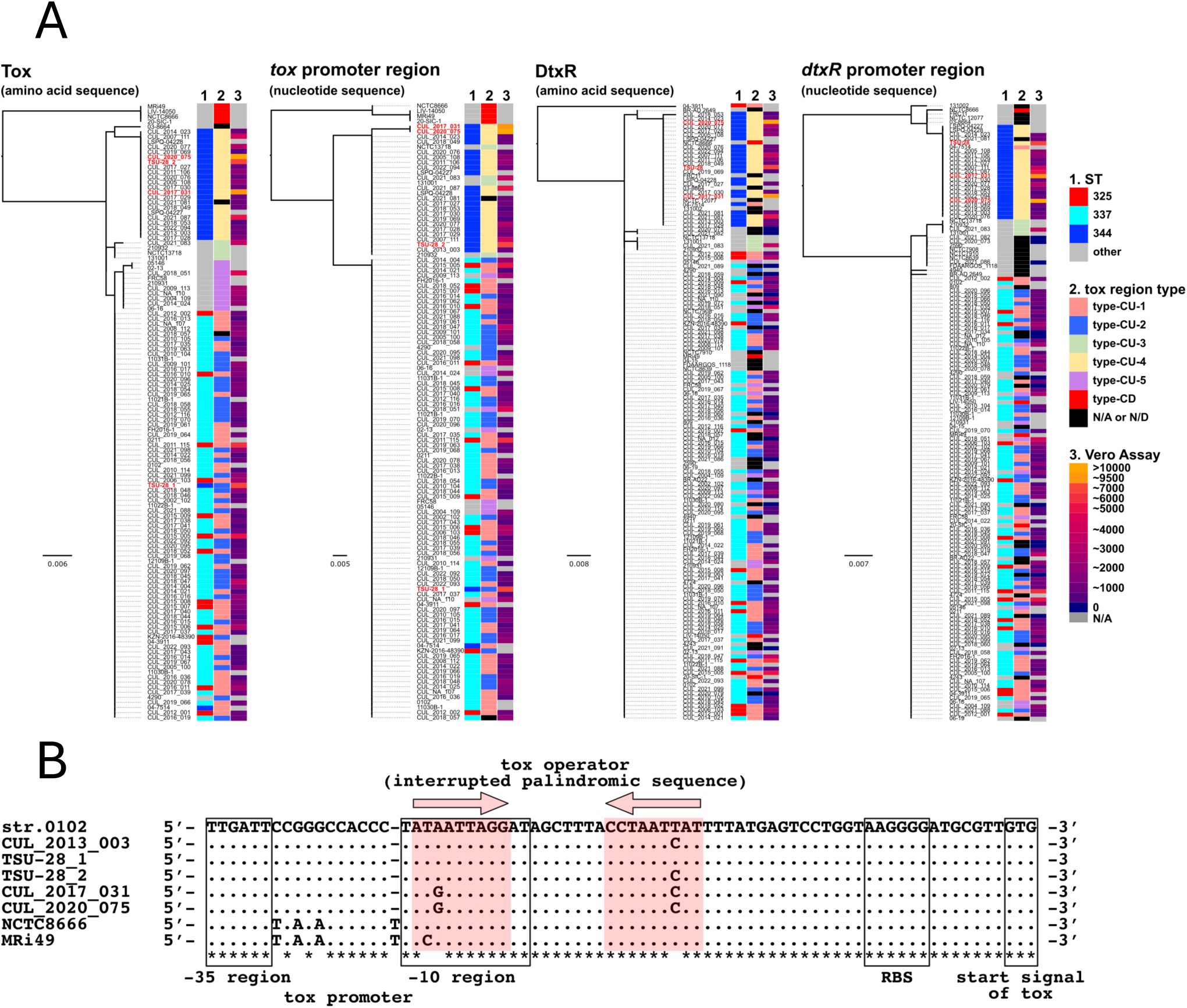
Phylogeny of tox-related sequences and the close-up view of the tox promoter region. The sequences of the toxicity-related genes and regions were analyzed. **A**. The *tox* and *dtxR* genes, along with their promoter-operator regions, were obtained from 106 isolates and 44 genomic sequences available in GenBank in March 2023 and compared; the resulting data were used to construct a phylogenetic tree with metadata. **B**. Nucleotide sequences of the operator-promoter region of the tox gene in ST325 outlying isolates (high producers). Dots indicate common nucleotides. Letters represent nucleotides that are different from common sequences.

## Discussion

In developed countries, *C. ulcerans* has emerged as one of the major pathogens causing diphtheria, sometimes surpassing the classical diphtheria pathogen, *C. diphtheriae* in case numbers. In contrast to the exclusive human pathogen *C. diphtheriae*, *C. ulcerans* is a zoonotic pathogen with a host range spanning many mammalian species, including humans and cats, thus allowing companion animals to serve as sources of zoonotic transmission to humans. With the accumulation of clinical information and isolates, some members of *C. ulcerans* are now recognized as independent bacterial species, *Corynebacterium silvaticum* [15] and *C. ramonii* [63]. In the present study, we analyzed 122 clinical and animal isolates obtained from Japan, including *C. ramonii* (*C. silvaticum* was not included because it has not yet been reported in Japan). In addition, genome sequences of European isolates recently published by Crestani et al. [39] and MLST data uploaded to the Institut Pasteur *Corynebacterium* database were included in the analysis.

Among Japanese isolates, ST337 was the predominant type, comprising 69.0% of all Japanese *C. ulcerans* isolates (69 out of 100 *C. ulcerans*; see Table S1), followed by ST325 (12 isolates; 12.0%). The composition of STs among the local isolates, including these two, showed significantly different patterns from those observed in Europe (France, Belgium, Germany, and the United Kingdom) (Figure 2). The unique predominance of a single MLST (ST337) in Japan, which is rare in Europe, suggests that a geographic barrier contributed to the pathogen’s composition. ST325, the second-largest group in Japan, ranked first in France and second in Germany. Phylogenetically, as shown in Figure 1, the two STs (337 and 325) were closely related to each other and placed in neighboring clusters (phylogenetic tree and circle 4), as Crestani et al. pointed out that they are in the same sublineage [39]; however, these two STs were nevertheless distinguishable from each other. Other predominant STs in Europe are ST331 in France and Germany and ST339 in Belgium (Figure 2). They were not present among the Japanese isolates or among those analyzed in the present study. Interestingly, in the United Kingdom, ST distribution patterns are quite different. In the United Kingdom, the European majority strains, ST325 and ST339, have not yet been reported in the public database (https://bigsdb.pasteur.fr/diphtheria/). The reason for this remarkable difference is unknown; however, the canal may have been a geographic barrier. In addition, based on the same database, the absence of these typical European STs was observed among isolates from Canada and Brazil (in the above database: data not shown). Such geographically distinct distributions have also been reported for other bacterial species. *Mycobacterium tuberculosis* has also been intensively studied [66]. In addition, for *Staphylococcus aureus* [67] and *Pseudomonas aeruginosa* [68], a globally differential distribution has been reported.

From a domestic viewpoint, geographic variance has been observed, even inside Japan. Figure 3 shows the geographic distribution patterns of *C. ulcerans* and *C. ramonii* isolates from Japan. The domestically predominant ST337 was distributed from the central to western regions of the country but not in the northern regions. In contrast, ST325, one of the predominant STs in Europe, has a smaller population but is exclusively distributed in the central to northern regions of the country. Additionally, the distribution of ST344 (*C. ramonii*) was skewed towards the western and southern regions. The factors contributing to this diversity are unknown; however, the pattern may reflect the history of *C. ulcerans* introduction and spread in Japan. Chronologically, as shown in Figures 3B and 3C, there were approximately two peaks during 2008–2012 and 2016–2022. In the earlier peak, the majority of ST337 isolates were from the Kansai region (the mid-west region), in contrast to the later peak, which was mainly from the Kanto region (Fig. 3B). In the Kanto region if the pathogen was introduced where little experience of *C. ulcerans*, such a predominance could be explained by “founder effect” in which a specific genetical group becomes predominant [69], when introduced initially “as a pioneer.” Another explanation was the migration of (for example) apparently uninfected hosts into the region, but this was unlikely because most of the hosts from which *C. ulcerans* was isolated were companion cats, which would not have been highly migratory. As described above, there was an interval between the two peaks. The number increased and diminished during 2008–2012, with a larger peak appearing in the 2015–2019 period. This might reflect social factors that have increased the general recognition of *C. ulcerans* infections among Japanese people. In the past, the NIID and the Ministry of Health, Labour and Welfare of Japan (MHLW) had started a campaign, at that time not huge, with official alerts on the MHLW website, directed at clinical veterinarians to promote the recognition of *C. ulcerans* infection as a zoonosis, first in 2002 and then updated in 2009. This may have been partially effective, leading to the first peak. In 2018, the alert was updated to mention a fatal case from 2016. Alerts were picked up by the media, including newspapers and TV [70]. Finally, the Chief Secretary mentioned it at a press conference because of the excessive “heat-up” [71]. This affected the recognition of *C. ulcerans*, especially among clinical veterinarians, more than any previous alert. Therefore, the 2016-2019 peak should be carefully observed with a background that probably indicates a bias on the clinical side.

Among the predominant ST 337 isolates (n=69) in Japan, 46 (66.7%) were from cats, and 22 (31.9%) were from humans. Among the isolates subjected to the present study, only one host was a dog. The reason for the high cat ratio among the isolates remains unclear. Except for the cat cases associated with five human cases (n=20), a large number of isolates from cats, especially during the 2016–2019 peak period, were sent to the NIID from commercial clinical testing laboratories that accept laboratory testing of swabs and isolates upon request by veterinarians, in most cases, private clinics. The remaining samples were obtained from prefectural and municipal laboratories. The high rate of cats might reflect a higher frequency of cat presentations to veterinarians, rather than a higher frequency of dog presentations. The number of dogs as companion animals in Japan is lower than that of cats, and continues to decrease [72]. However, this does not fully explain the large differences in the numbers of cat and dog isolates.

The toxigenicity studies employed a combination of *tox*-PCR and Vero cell cytotoxicity assays as routine tests. The benefit of the Vero cell test is its quantitative nature and high sensitivity, compared with the Elek test, which is less quantitative and sometimes not suitable for *C. ulcerans,* which generally produces less toxin than *C. diphtheriae* [73]. The Vero cell assay thus helps avoid the potential misjudgment of low-toxin producers as NTTB (non-toxigenic but toxin gene-bearing). In the present study, we reliably estimated and compared the toxin production of *C. ulcerans* isolates, even at low levels. However, in our toxin production estimate, there was a limitation: the toxin production was carried out under a specific fixed condition, namely, in one type of liquid medium (Elek broth [2]) at OD600=0.5. However, to ensure a reliable comparison of toxin production under these conditions, the liquid cultures of all the isolates were analyzed simultaneously at the mid-log phase and standardized to an optical density of 0.5 at OD500 to compensate for small differences. As a result, ST337 revealed to contain comparably low producers with less diversity than other ST groups, such as ST325 or ST332, although it included a fatal case [4]. Interestingly, among the ST337 isolates, a significant difference in toxin production (p<0.01) was observed between isolates with the CU-1 and CU-2 prophages (Figure 7). The two types differed in the attachment site of the prophage but were highly homologous to each other (Figure 4). As shown in Figure 8, no significant differences were observed between the two types in the nucleotide sequences of the tox gene, the dtxR regulatory gene, and their respective promoter sequences. In contrast, the ST325 and ST331 isolates produced larger amounts and more diverse toxins. Remarkably, ST344, representing *C. ramonii*, was much more diverse in terms of toxin production. The typical outliers among the ST344 isolates, CUL-2017_031 and CUL_2020_075, showed approximately 15- and 30-fold higher levels of toxin production, respectively, than the average level of ST337 isolates (see Fig. 7 and Table S1). These levels were equivalent to or greater than those of the two *tox* gene-bearing TSU-28 strains [65] (Table S1). High producers CUL-2017_031 and CUL_2020_075 harbored nucleotide substitutions in the tox gene operator region. (Figure 8B). The substitutions were located within the palindromic structure of the promoter region, partially overlapping the -10 region [74]. The palindromic sequence has previously been shown to be crucial for regulating the transcription of the tox gene, not by forming a stem-loop structure, but by being recognized by a dimer of the repressor (the *dtxR* gene product) [75–77]. Nucleotide substitutions affecting transcription were found within the palindromic sequence, but in the very center between 5’-TTAGG and CCTAA-3, where the substitutions in CUL-2017_031 and CUL_2020_075 were not located. Nevertheless, there may be unknown mechanisms underlying the apparently different characteristics of toxin production, related to the nucleotide sequences of the above regions. As shown in Figure 8A, separate clades were formed between ST337 and ST344 (*C. ramonii*) based on phylogenetic analysis of the amino acid sequences of the *tox* and *dtxR* gene products and the nucleotide sequences of their promoter regions.

Regarding nontoxigenic isolates, Crestani et al. [39] pointed out that clonal group 339 (corresponding to ST339) is the most common among European *tox* (-) isolates, whereas in our Japanese *C. ulcerans* collection, ST339 was absent not only among the tox(-) isolates, but was also completely absent. Instead, among our collection, 13 isolates were tox(-), including 337 (n=5), 699 (n=2), 327 (n=2), 811 (n=1), 971 (n=1), and 1179 (n=2).

Our study demonstrates that multiple types of *tox*-containing prophages and chromosomal regions exist among *C. ulcerans* Japanese and global isolates. The most prominent types in Japan were CU-1 and CU-2 (Fig. 1; see also Table S1), which were found in the most common ST337 isolates. The nucleotide sequences of these intact prophages showed approximately 99% identity; the only notable difference was in their integration sites within the chromosome. This could have resulted from prophage exchange events by the induction and re-infection of bacteriophages among ST337 organisms. Notably, the CU-1 and CU- 2 prophage types were found almost exclusively in ST337 among the Japanese isolates sequenced in the present study, with 12 exceptions of the ST325-CU-1 type. In contrast, in Europe [39] ST325 is the only host accommodating the CU-1 type prophage. The CU-2 type was, except the ST337, only found in five ST1011 isolates [78] in Japan. In addition, there was an ST344 double lysogen with CU-2 and CU-4 (PAI) type prophages [65], which is the only case reported in Japan. However, a large number of double lysogens with CU-2 and CU-4 (PAI) type prophages have been reported in Europe [39].

Three incomplete types of prophages exist, which can also be regarded as PAIs, as described by Meinel et al. [38] and Crestani et al. [39]. They are designated as CU-3, 4, and 5 in Figure 4. The CU-3 type contained multiple reading frames that represented *C. ulcerans*-type phage components (Figure 4, CU-3, blue ORFs, and yellow bar beneath). In the CU-4 type, which was exclusively associated with ST344 (*C. ramoni*) when found alone or with a CU-2 type prophage, a suspected trace of a *C. ulcerans*-type prophage was not present, but another suspected trace of a prophage of *C. diphtheriae*-type appeared (blue bar). The possible phage trace structure was conserved as the CU-5 type, ultimately corresponding to the uppermost *tox*(-) scheme (*C. ulcerans* NCTC7910).

On the opposite (lower) side of Figure 4 (CD types), schemes for isolates not found in Japan are shown. These represent European isolates in which the nucleotide sequence of the prophage regions is non-homologous to *C. ulcerans* types (CU-1 and 2), but homologous to that of *C. diphtheriae* (see the lowermost scheme in Figure 4 representing *C. diphtheriae* NCTC13129, the first opened genome). Nevertheless, in the schemes of all types, the surrounding regions were highly homologous to each other, including those of the CU types, suggesting that any type of *tox*-containing prophage prefers two attachment sites on the chromosome, both of which are located very close to each other. The presence of two attachment sites in the *C. ulcerans* chromosome was reported by Seto et al. [20] in 2008.

There are two “hybrid-type” prophages, CU-6 and CD-5, among genomes of isolates outside Japan (Figure 5). Type CU-6 could be considered a chimera between the CU-1 and CD-1 types (middle to lower part of Figure 5). In the nucleotide sequence of the CD-5 type prophage, a region homologous to a *C. diphtheriae* protein unrelated to the *tox* gene (the upper part of the Figure) was observed, suggesting a history of insertion of the *tox* gene and its neighboring nucleotide sequences into such an unrelated prophage. These incomplete types can be regarded as PAIs. The animal sources of the *C. ulcerans* isolates analyzed in the present study were diverse, but most were cats. Unfortunately, we could not find a relationship between the animal of origin and MLST or prophage type.

In contrast, toxin production was partly associated with MLST type (Figure 7) but was not explained by the nucleotide sequences of the upstream regions of the tox and dtxR genes (Figure 8).

This study revealed that the local specificity of *C. ulcerans* isolates differed between Japan and Europe. Notably, the majority of isolates belonged to ST337, which is rare in Europe, suggesting a different route of introduction of C. ulcerans into Japan and Europe. The difference was not only in MLST types but also in prophage types, revealing that prophage types were closely associated with MLST types.

*C. ulcerans*, including *C. ramonii* and *C. silvaticum*, are emerging pathogens that override *C. diphtheriae* in developed countries. Diphtheria caused by *C. diphtheriae* is also re-emerging in Africa, resulting in WHO alerts [79], and in Europe among migrants and asylum seekers [80]. Although the route of infection differs for the zoonotic pathogen C. ulcerans, an intensive public health approach is equally important for diphtheria-causing pathogens. Recognition of the disease, especially when aided by the media, is very important for clarifying the pathogen’s prevalence. Increased recognition has also helped identify human cases that might otherwise have gone unrecognized without more careful observation by clinicians and clinical veterinarians.

## Materials and Methods

### Bacterial strains and online data

*C. ulcerans* strains used in this study and their metadata are listed in Table S1. A total of 106 Japanese isolates were sequenced in the present study and all of them were subjected to the toxin production assay. Publicly available *C. ulcerans* genome sequences obtained for analysis were also analyzed, mainly those reported by Crestani et al. [39]. In addition, MLST data were obtained from the Institut Pasteur database (https://bigsdb.pasteur.fr/diphtheria/), and the entire dataset is presented in Table S1. The Japanese isolates were found in humans, cats, dogs, and other animals, as well as the environment where the patient and suspected animals resided. Most human isolates are sent to the National Institute of Infectious Diseases (NIID) after isolation at hospitals where patients are treated. Animal isolates recovered from veterinary clinics or clinical testing laboratories were sent to the NIID. Some animal isolates were obtained by active surveillance of companion animals associated with human cases or wildlife. All isolates were stored at -80 °C in 50 % glycerol until culturing.

### NGS genome sequencing

Isolates to be sequenced were reconstituted on sheep blood agar plates (Nissui Seiyaku, Tokyo, Japan, cat# 50001) at 37 °C for 24–48 h. The reconstituted bacterial cells (one loopful on a 1-uL disposable loop) were inoculated into brain-heart infusion liquid medium (BHI: Becton-Dickinson, Bacto Brain-Heart Infusion) and were cultivated as a preculture at 37 °C with shaking. Then, 100 ul of the preculture was transferred to fresh BHI and similarly cultured. After 24 h of cultivation, the bacterial cells were collected by centrifugation (3500 rpm, 5 min). The optical density of the culture was measured at 600 nm prior to centrifugation using a GeneQuant Pro spectrophotometer (Pharmacia, GE Healthcare, Tokyo, Japan). The cell pellet was resuspended in phenol:chloroform: isoamyl alcohol (25:24:1). The bacterial cell suspension was kept at 4 °C until use. Bacterial cells were disrupted by bead-beating using a Geno Grinder at 1,500 rpm for 10 min. Genomic DNA was extracted using a bead-beating mixture and purified with a ZR-96 Zymoclean Gel DNA Recovery Kit (Zymo Research). DNA-seq libraries were constructed using a QIAseq FX DNA Library Kit (QIAGEN, Tokyo, Japan). DNA sequencing was performed on a NextSeq 500 platform using the NextSeq 500/550 Mid Output Kit v2.5 (150 bp paired-end; 300 cycles).

### Estimation of toxin production by Vero cell cytotoxicity test

Bacterial isolates were reconstituted from frozen stock (see "*Bacterial strains and online data*") on sheep blood agar plates. The reconstituted cells (one loopful on a 1-uL disposable loop) were seeded in 2 mL of Elek broth [2] and incubated at 37 °C with shaking, as precultures. After 24 h of preculturing, 100 µL of the preculture was then seeded into 2 mL of fresh Elek broth and incubated at 37 °C for 48 h with shaking. After 48 h of incubation, the cells were removed from the incubator. Then, 200 ul of culture was recovered before centrifugation and diluted 5-fold with sterile water to measure the optical density at 600 nm. Elek broth diluted five-fold was used for background subtraction. The remaining culture was centrifuged at 3500 rpm for 5 min to precipitate the bacterial cells. The supernatant was sterilized by filtration through a 0.22 um membrane filter and stored at 4°C until use. Vero cell cytotoxicity tests were performed as previously described by Miyamura et al. [44, 45]. Briefly, 25 uL of serial twofold dilutions of bacterial culture filtrate, 75 ul of minimal essential medium (MEM) containing 3% of fetal calf serum, and 50 ul of Vero cell suspension (3 × 105/mL in MEM) were added. After 4 d of incubation at 37 °C in a 5% CO2 atmosphere in a humidified incubator, the endpoint (50% cytotoxicity) was determined by visual inspection with a Cell3iMager Duos image analyzer (SCREEN holdings, Shiga, Japan) and confirmed by naked-eye microscopic observation. The observed cytotoxicity was confirmed to be due to the diphtheria toxin by neutralization with a diphtheria antitoxin (Japanese National Standard Diphtheria Antitoxin lot 10).

### Draft genome sequence analysis

Adapter trimming of short reads was performed using fastq-mcf version 1.04.636 (https://openbioinformaticsjournal.com/VOLUME/7/PAGE/1/) with default parameters, followed by low-quality trimming using Sickle version 1.33 (https://github.com/najoshi/sickle) with the following parameters: minimum length threshold ‘-l 40.’ The draft genome sequence was assembled using SKESA version 2.3.0 [46] with Illumina short-read data only. Gene annotation was performed using DFAST version 1.2.3 [47] with the DFAST default database and Virulence Factors Database [48]. Multilocus sequence typing (MLST) was performed using the “mlst” program version 2.16.2 (Seemann T, mlst Github https://github.com/tseemann/mlst (accessed on August 29, 2022)) with the PubMLST database (https://pubmlst.org/ (accessed on August 29, 2022)).

### SNV Phylogenetic Analysis and Pan-Genome Analysis

Deposited genome assemblies and/or Illumina raw sequence reads were retrieved from the NCBI for the Biotechnology Information database (Table S1). For SNV analysis, the complete genome sequence of *C. ulcerans* 0102 (AP012284.1) was selected as the reference sequence. The repeat and prophage regions of the reference sequences were analyzed using NUCmer (MUMmer 3.0) [49] and PhiSpy version 4.2.21 [50], respectively. If the available data consisted only of contig sequences from the NCBI database, Ngsngs version 0.9.2 [51] was used to simulate 150-mer paired-end reads with a 200 bp insert length. Read mapping was performed using BWA-MEM [52] version 0.7.17, with default parameters against reference chromosomal sequences. The mapping coverage and statistics were calculated using BEDTools version 2.31.1 [53] and SAMtools version 1.20 [54], respectively. Duplicate reads were marked using the MarkDuplicates tool in GAKT version 4.5.0.0 [55], followed by variant calling using the Haplotype Caller tool in GAKT. Single nucleotide variants (SNVs) and insertions/deletions (indels) were extracted using the SelectVariants tool from GATK, and filtered with the VariantFiltration tool using the following parameters: for SNVs, QD < 2.0, QUAL < 30.0, SOR > 4.0, FS > 60.0, MQ < 20.0, MQRankSum < –12.5, isHet == 1, ReadPosRankSum < –8.0, DP < max(mean depth / 10, 5); for indels, QD < 2.0, QUAL < 30.0, SOR > 10.0, FS > 200.0, isHet == 1, ReadPosRankSum < –20.0, DP < max(mean depth / 10, 5). To avoid excessive filtering in low-coverage regions, the depth (DP) threshold was set to the greater of the mean depth divided by 10 or 5 (i.e., DP < max(mean depth / 10, 5)). Base quality scores were recalibrated using the BaseRecalibrator and ApplyBQSR tools from GATK, followed by final variant calling and filtering using a previously described method. To predict recombination regions, consensus sequences were constructed from the SNV data and reference sequences using BCFtools version 1.21 [54] and BEDTools. All SNV datasets were merged using vcftools version 0.1.16 [56], followed by filtering of SNVs located in non-core, repeat, and predicted prophage regions. SNVs within repeat regions and predicted prophage regions were removed, and recombination regions were predicted using Gubbins version 3.4 [57]. Core genome SNV phylogenetic analysis was performed by IQ-TREE version 2.4.0 [58] with the parameters ‘-m MFP -b 100’ [59], followed by visualization using Fandango version 1.3.0 [60] and interactive tree of life (iTOL) version 3 [61]. SNV network analysis was performed using the median-joining network method in PopART (http://popart.otago.ac.nz (accessed August 29, 2022)). Pan-genome analysis of predicted open reading frames (ORFs) was performed using Roary version 3.12.0 with default parameters [62].

## Acknowledgments

We are grateful to Atsuko Aoki, Mayumi Arituka, Shouji Asakura, Fujifilm Vet Systems, Koji Fujimoto, Noriko Hagiwara, Toshinaga Hasegawa, Kaoru Hatakeyama, Hideki Hino, Sho Hirai, Ryuichi Ikemoto, Asuka Inomata, Kazuhiko Ishikawa, Ryoko Jikihara, Hideaki Kariya, Hoken Kagaku, Inc., Chihiro Katsukawa, Yuta Kawakami, Jun Kawase, Shun Kikuchi, Takahiro Kinebuchi, Marina Kon, Tomomi Kono, Eriko Koyama, Miroku Medical Laboratory, Yoshiteru Murata, Hisami Nagami, Hiroshi Nakajima, Fumihiro Ochi, Kikuyo Ogata, Masao Ogawa, Norio Okazaki, Tomotake Sakai, Tadashi Sankai, Junji Seto, Yukiji Seto (Shirai), Miwako Shichikum, Keiji Shimada, Ikuyo Shimono, Kunihiro Shono, Mutsumi Tanimura, Kensuke Tsuneyasu, Yoshie Tsunomori, Kaoru Umeda, Masato Wakamatsu, Yuko Watanabe, Motohiko Yamada (in alphabetical order) for the isolates and cooperation. We also thank the physicians and veterinarians who contributed to this project for their support throughout this study.

## Funding

This work was supported by H19-Shinkou-Ippan-009, H22-Shinkou-Ippan-010, JP15fk0108005, JP16fk0108117, JP17fk0108217, JP18fk0108017, JP19fk0108097, JP20fk0108097 and JP21fk0108101.

No open access license has been selected.

## Data availability

Nucleotide sequence data are publicly available with accession numbers listed in Supplementary Table S1.

