## Supplemental Figures 1-4 for "Comparative genome analysis of *Corynebacterium ulcerans* Japanese isolates revealed domestically and globally diverse geographical distribution of the organism of different types"

[Figure S1]

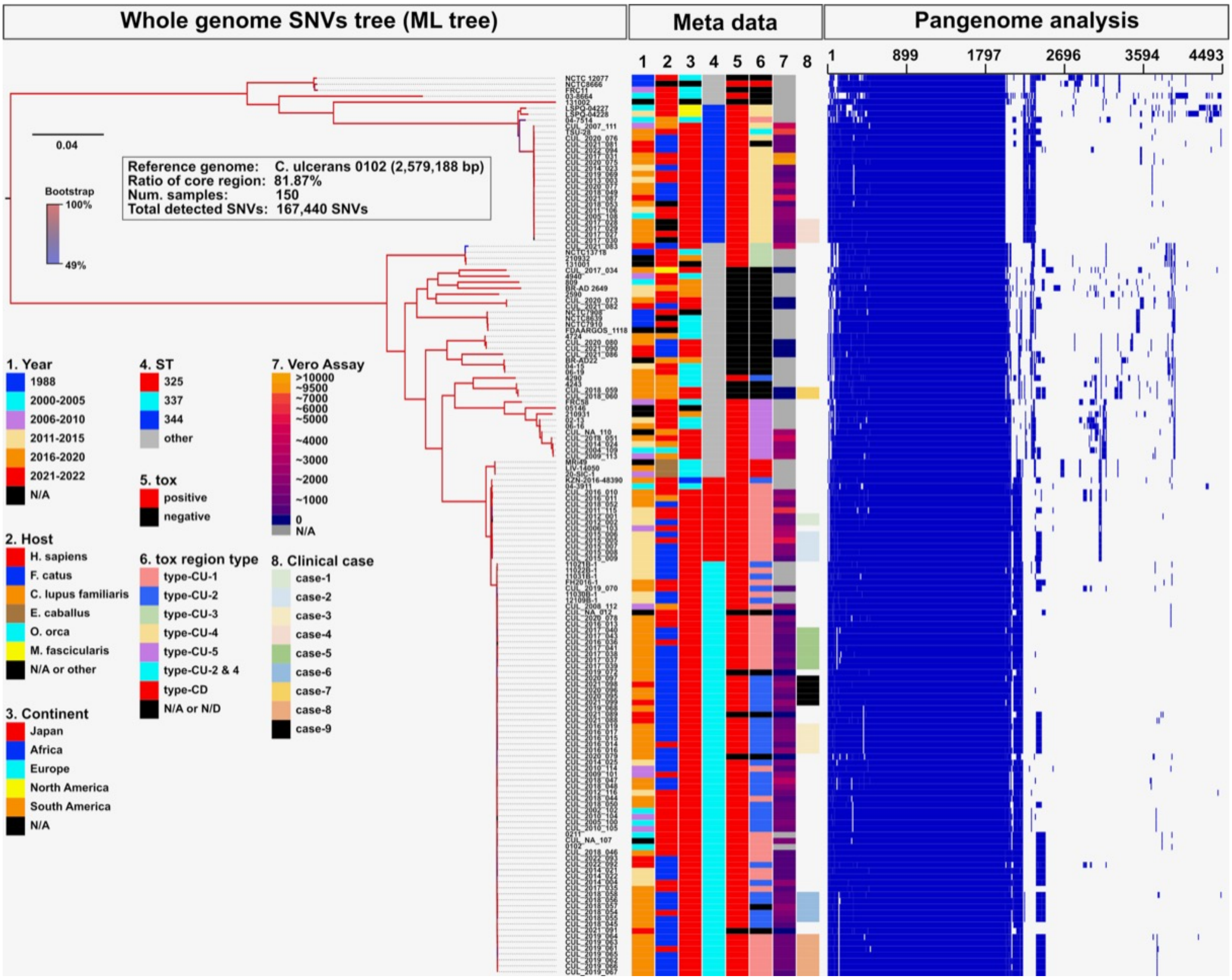


*Figure S1. Phylogenetic tree and metadata of genomes of 150 C. ulcerans and C. ramonii isolates.* Genome sequences obtained in the present study (n=106) and those available in March 2023 (n=44) were analyzed as described in Materials and Methods and expressed in phylogenetic tree. Metadata associated to each isolate and those obtained from database are also shown (bars 1-3 and 8). MLST types (as ST), presence/absence of the *tox* gene, prophage types (as *tox* region type) are shown in bars 4 to 6, respectively. The bar 7 indicates the level of toxin production in Elek broth at OD=0.5, measured by Vero cell assay. Blue and blank areas in "Pangenome" represents represent the presence and absence of the corresponding genomic regions, respectively.

[Figure S2]

Geographic distribution of animals of origin.

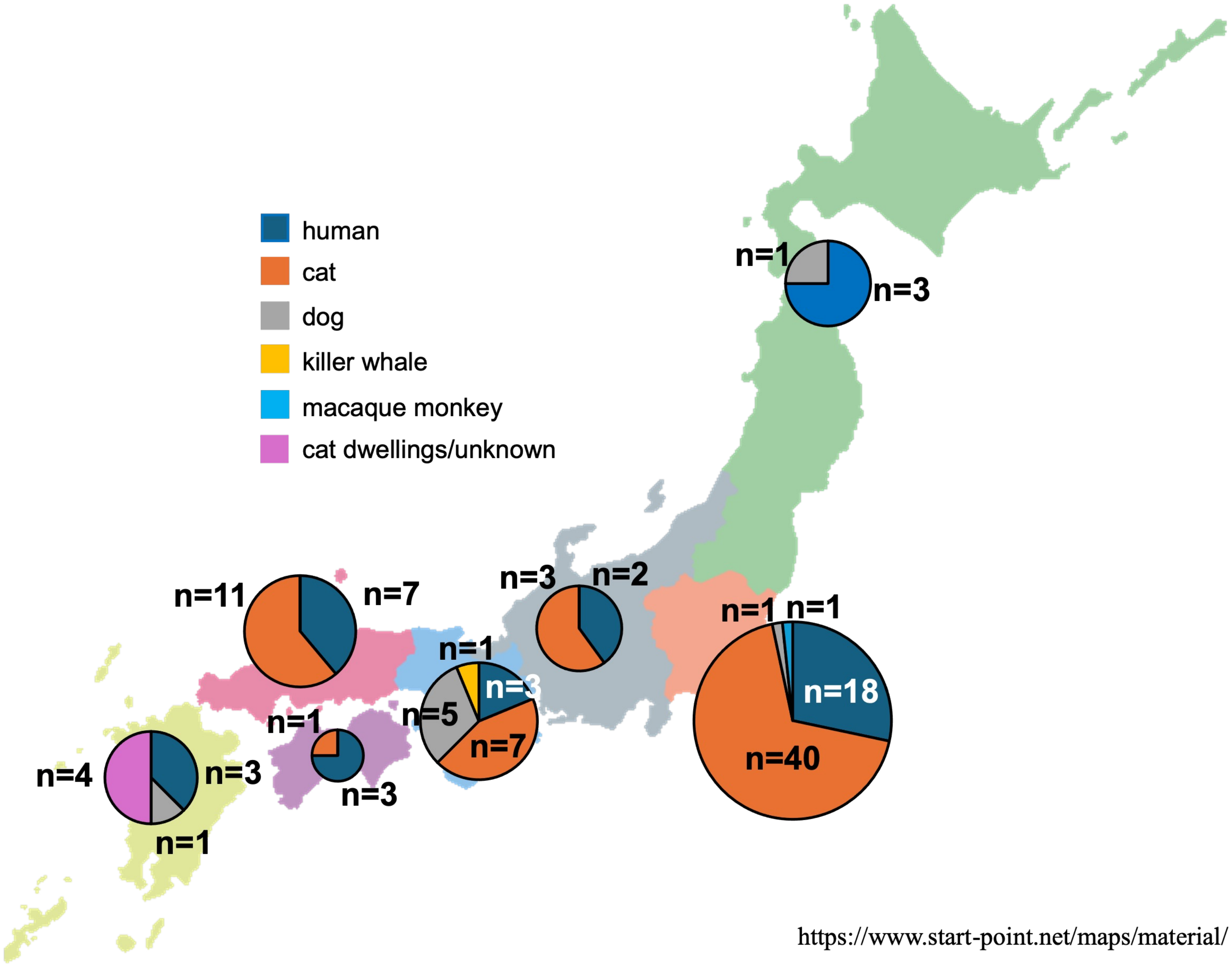

### [Figure S3]

Geographic distribution of prophage types.

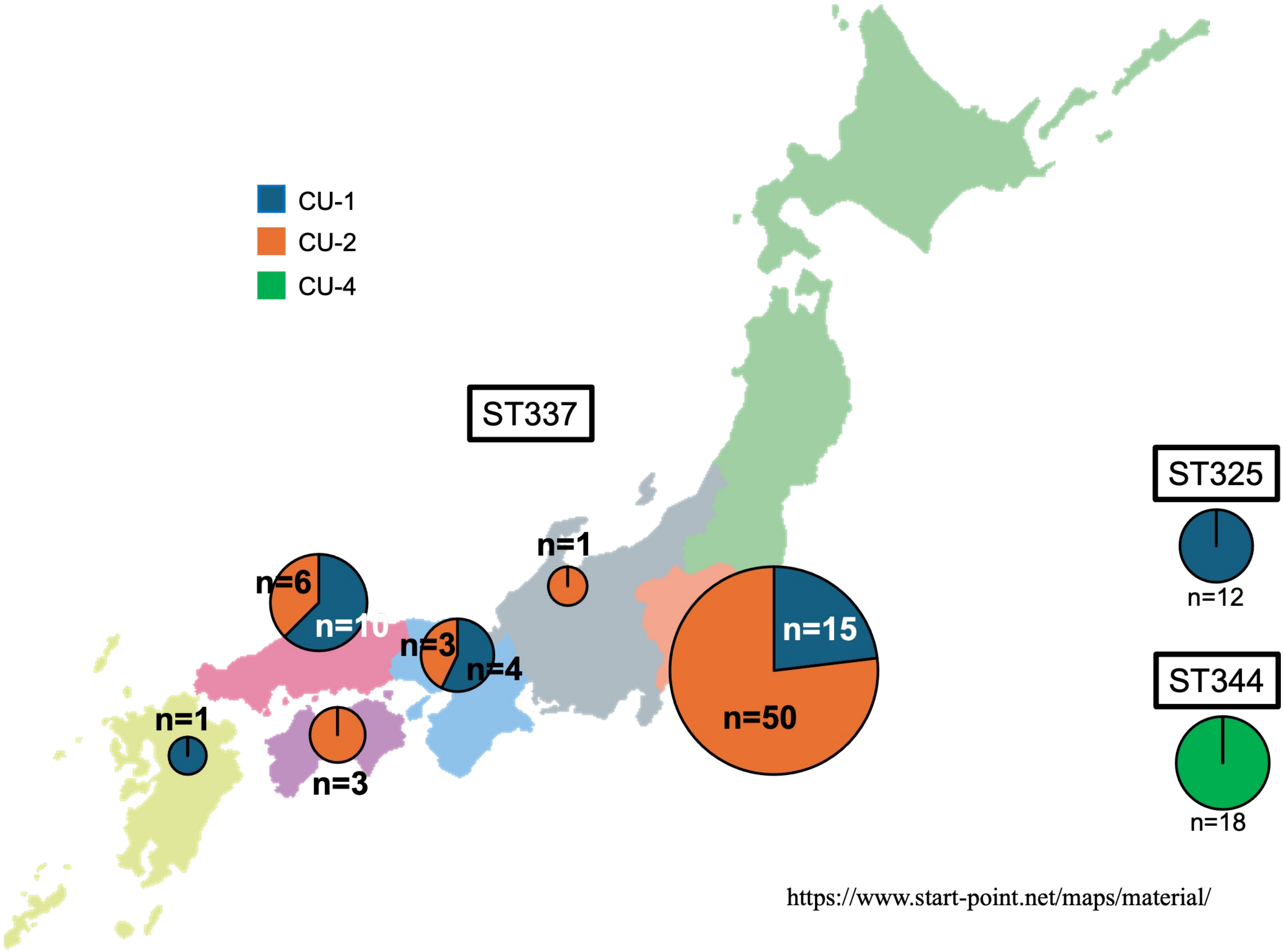

[Figure S4]

Geographic distribution of clinical cases associated with animals.

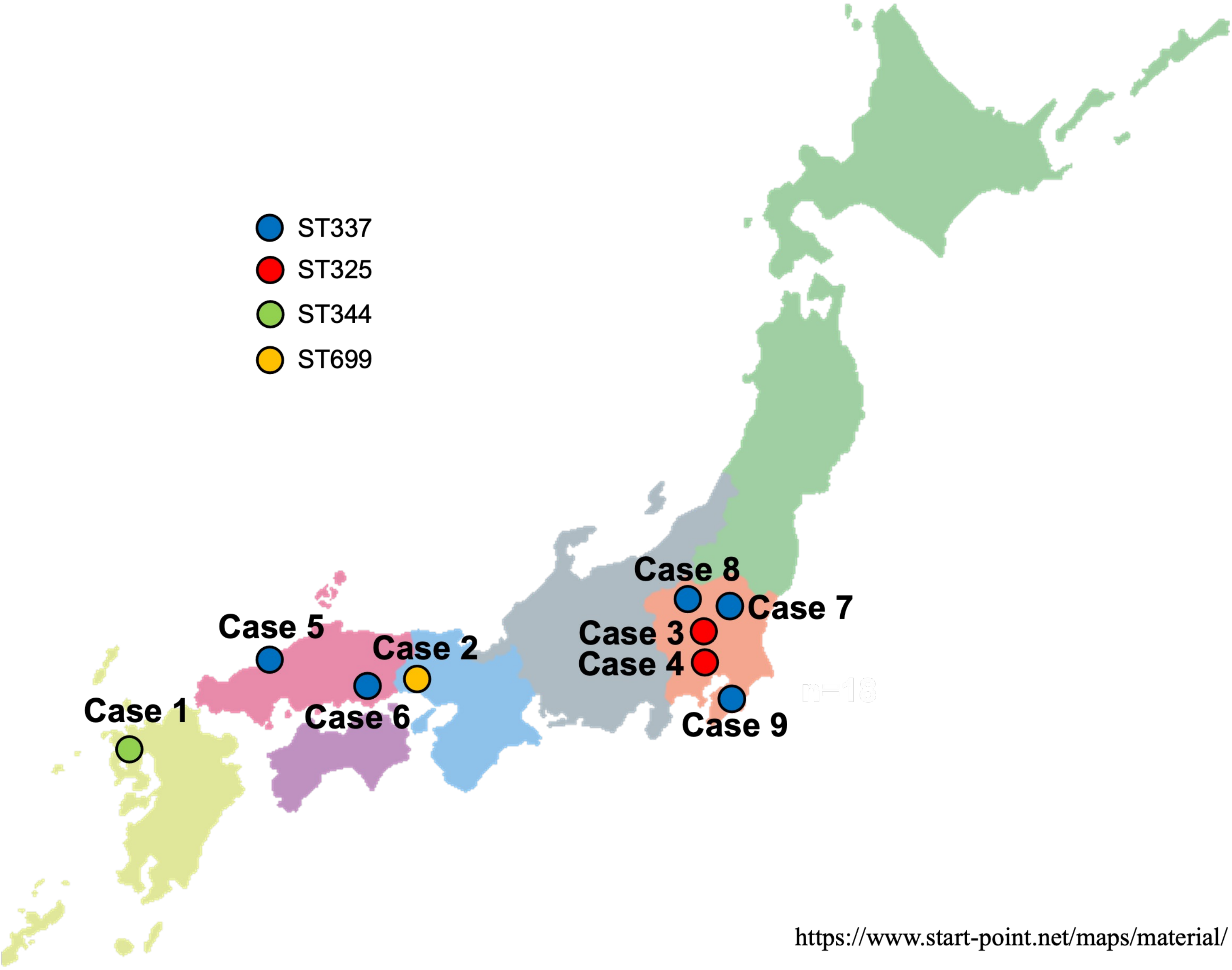
